# Low-dose doxorubicin drives caveolin-1 depended re-epithelialization of breast cancer cells as a mechanism of cancer plasticity

**DOI:** 10.64898/2026.09.28.754929

**Authors:** Alexandros Damalas, Marijonas Tutkus, Ioannis D Kyriazis, Zozo Outskouni, Ernesta Vitulskienė, Panteleimon M Dimovelis, Soultana Kafetzi, Charlampos Angelidis, Panagiotis Liakos, Varvara Trachana

## Abstract

Breast cancer progression is driven by dynamic changes in epithelial plasticity, membrane organization, and intracellular signaling, yet the effects of sustained low-dose chemotherapy on these processes remain poorly understood. Here, we investigated the impact of prolonged low-dose doxorubicin on membrane remodeling, epithelial phenotype, membrane-associated Ras lipid-anchor localization, and autophagy in mesenchymal-like MDA-MB-231 breast cancer cells. Low-dose doxorubicin significantly increased Caveolin-1 expression and enhanced E-cadherin protein levels, accompanied by a transition toward a more compact epithelial-like morphology with increased cell-cell contacts. Live-cell imaging demonstrated a significant reduction in the membrane-to-cytoplasm fluorescence ratio of the lipid-anchored GFP-tH probe, indicating redistribution from the plasma membrane to the cytoplasm following treatment. Analysis of autophagy-related proteins revealed decreased LC3-I together with increased LC3-II, ATG5, and p62 expression, consistent with autophagosome accumulation and impaired autophagic flux. Collectively, our findings demonstrate that low-dose doxorubicin promotes extensive remodeling of plasma membrane organization, epithelial plasticity, membrane-associated lipid-anchor localization, and autophagy. This integrated response reveals previously unrecognized links between membrane architecture, Ras membrane association, and autophagy during phenotypic reprogramming of breast cancer cells, providing mechanistic insight into cellular adaptations elicited by sub-cytotoxic doxorubicin exposure.

## Introduction

Breast cancer remains the most frequently diagnosed malignancy among women worldwide and continues to represent one of the leading causes of cancer-related mortality (Lukasiewicz et al., 2021). Although major advances in early diagnosis, molecular classification, and targeted therapies have significantly improved patient survival, metastatic disease remains largely incurable (Nicolini et al., 2025). The ability of tumor cells to survive therapeutic pressure and subsequently acquire more aggressive phenotypes has emerged as one of the principal barriers to successful long-term treatment. Increasing evidence indicates that these adaptive responses are driven not only by genetic alterations but also by profound remodeling of membrane organization, intracellular signaling, cytoskeletal architecture, protein trafficking, and stress-response pathways (Lemanowicz et al., 2026). Understanding how these interconnected processes enable cancer cells to survive chemotherapy has become an important objective in modern cancer biology (Dhiman et al., 2026).

Among the mechanisms underlying tumor adaptation, epithelial–mesenchymal transition (EMT) and the reverse process, mesenchymal–epithelial transition (MET), represent fundamental determinants of cancer progression (Lamouille et al., 2014). EMT promotes migration, invasion, stemness, immune evasion, and therapeutic resistance, whereas MET facilitates re-establishment of epithelial characteristics that support metastatic colonization and survival (Allgayer et al., 2025). Rather than binary events, EMT and MET are now recognized as dynamic and reversible processes involving hybrid intermediate phenotypes regulated by extracellular signals, plasma membrane organization, cytoskeletal remodeling, and intracellular trafficking (Damalas et al., 2026, Datta et al., 2021). Mesenchymal-like breast cancer cells therefore possess remarkable phenotypic plasticity that enables continuous adaptation to changing microenvironmental and therapeutic conditions (Luo et al., 2015).

Triple-negative breast cancer (TNBC) is among the most aggressive breast cancer subtypes because of its high metastatic potential, molecular heterogeneity, and lack of targeted therapeutic options (Newton et al., 2022). MDA-MB-231 cells are widely used as a representative mesenchymal-like TNBC model characterized by reduced E-cadherin expression, enhanced migratory behavior, and constitutive activation of oncogenic signaling pathways (Liu et al., 2021).

Doxorubicin remains a cornerstone of breast cancer chemotherapy (Johnson-Arbor et al., 2026). At conventional concentrations, it induces DNA damage, oxidative stress, and apoptosis (Pilco-Ferreto and Calaf, 2016). However, accumulating evidence demonstrates that prolonged exposure to sub-cytotoxic concentrations activates adaptive programs rather than extensive cell death (Kciuk et al., 2023). Low-dose chemotherapy can remodel plasma membrane organization, alter oxidative signaling, regulate protein trafficking, influence autophagy, and modify cell adhesion (Chan et al., 2016). These adaptive responses may ultimately determine whether tumor cells undergo elimination or survive treatment. Nevertheless, the molecular pathways integrating these responses remain incompletely understood.

The plasma membrane has emerged as an active signaling platform rather than a passive structural barrier. Caveolae and their principal structural protein Caveolin-1 organize membrane nanodomains and regulate lipid homeostasis, mechano-transduction, endocytosis, and compartmentalization of signaling complexes (Filippini and D’Alessio, 2020, Parton and Simons, 2007). Caveolin-1 has context-dependent functions in cancer, acting either as a tumor suppressor or promoter depending upon tumor stage and cellular context (Quest et al., 2008). Importantly, Caveolin-1 has also been implicated in stabilization of adherens junctions, regulation of E-cadherin (Katsuno-Kambe et al., 2021), membrane mechanics, and Ras signaling, suggesting that it may coordinate multiple adaptive pathways simultaneously (Damalas et al., 2022, Abankwa and Gorfe, 2020, Ariotti et al., 2014, Ariotti et al., 2015).

E-cadherin is the principal epithelial adhesion molecule responsible for adherens junction assembly and tissue integrity (Coopman and Djiane, 2016). Loss of E-cadherin is a defining hallmark of EMT and correlates with invasion, metastasis, and poor prognosis, whereas restoration of E-cadherin accompanies MET (Bruner and Derksen, 2018). The relationship between Caveolin-1 induction and E-cadherin restoration during chronic exposure to low-dose chemotherapy remains poorly characterized and may reveal mechanisms underlying adaptive epithelial reprogramming.

Membrane architecture also determines the spatial organization of Ras proteins. H-Ras requires post-translational lipid modification for stable association with the plasma membrane, and its signaling activity depends upon correct membrane compartmentalization. Alterations in lipid composition, membrane curvature and caveolar organization can redistribute H-Ras from the plasma membrane to intracellular compartments, thereby modifying downstream MAPK and PI3K signaling (Damalas et al., 2026, Damalas et al., 2022) (Liang et al., 2019) (Zhou and Hancock, 2023, Zhou et al., 2017, Zhou and Hancock, 2018). Consequently, changes in localization of the H-Ras tH lipid anchor represent a sensitive indicator of membrane remodeling during therapeutic adaptation.

Autophagy constitutes another major adaptive response activated during cellular stress. Emerging evidence further suggests extensive crosstalk between Caveolin-1 and autophagy, indicating that membrane organization may directly regulate autophagic homeostasis (Yang et al., 2025).

Although EMT, Caveolin-1 biology, Ras compartmentalization, and autophagy have each been investigated extensively, these pathways have generally been studied independently. Their coordinated regulation during chronic exposure to low-dose doxorubicin remains largely unexplored. Consequently, whether plasma membrane remodeling functions as the central event integrating epithelial plasticity, Ras localization, and autophagic adaptation has remained an important unanswered question.

Our previous studies proposed that membrane remodeling and membrane curvature actively regulate oncogenic signaling and epithelial plasticity (Damalas et al., 2022). The present work extends these concepts experimentally by investigating how chronic exposure to 12.5 nM doxorubicin remodels membrane organization in mesenchymal-like MDA-MB-231 cells. Unlike previous studies examining isolated pathways, this study simultaneously evaluates Caveolin-1 expression, E-cadherin restoration, and redistribution of the H-Ras tH (Damalas et al., 2026) experimental model.

The novelty of this study lies in demonstrating that prolonged low-dose doxorubicin induces a coordinated adaptive remodeling program characterized by Caveolin-1 upregulation, partial mesenchymal-to-epithelial transition, redistribution of the H-Ras lipid anchor from the plasma membrane to the cytoplasm, and dysregulated autophagic flux. To our knowledge, this is the first study integrating these biological processes into a unified mechanistic framework of chemotherapy adaptation in breast cancer cells.

Collectively, the present findings support the concept that plasma membrane remodeling is not simply a consequence of chemotherapeutic stress but represents an upstream regulatory mechanism coordinating epithelial differentiation, Ras spatial organization, and autophagic adaptation. By integrating biochemical analyses, live-cell imaging, and protein localization studies, this work provides a comprehensive mechanistic framework explaining how chronic low-dose chemotherapy reshapes tumor cell biology beyond classical cytotoxicity. These findings identify membrane remodeling as a potential therapeutic vulnerability and establish a conceptual basis for future strategies aimed at limiting tumor adaptation and therapeutic resistance.

## Materials and Methods

### Cell culture and doxorubicin treatment

Cells were cultured in Dulbecco’s Modified Eagle Medium (DMEM; Gibco, Thermo Fisher Scientific) supplemented with 10% fetal bovine serum (FBS), 100 U/mL penicillin, and 100 μg/mL streptomycin at 37 °C in a humidified incubator containing 5% CO₂. For all experiments, cells were seeded and allowed to adhere for 24 h. The culture medium was then replaced with DMEM containing 5% FBS, and cells were treated with 12.5 nM doxorubicin for 24 h. Untreated control cells received an equivalent volume of vehicle under identical culture conditions. The MDA-MB-231 cell line was kindly provided as a gift from Professor Moshe Oren.

### Live-cell imaging

For live-cell imaging experiments, 18,000 cells were seeded in μ-Dish 35 mm glass-bottom imaging dishes (ibidi GmbH, Germany). After 24 h, cells were transiently transfected with 100 ng of the GFP–tH plasmid using Lipofectamine 2000 (Thermo Fisher Scientific) according to the manufacturer’s protocol. Twenty-four hours after transfection, the culture medium was replaced with DMEM containing 5% FBS, and cells were treated with 12.5 nM doxorubicin for an additional 24 h. Live-cell imaging was performed using a Zeiss inverted laser-scanning confocal microscope (Zeiss laser scanning microscope 800, Zeiss LSM 800, Zeiss Germany) equipped with Zeiss a C-Apochromat 40x/1.2 W Corr objective len. GFP fluorescence was visualized using the appropriate filter set, and images were acquired using identical microscope settings for all experimental groups. Image acquisition and analysis were performed using Zeiss ZEN software.

The image acquisition and quantitative analysis strategy followed the methodology previously described by Damalas et al. (Scientific Reports) for analysing Ras membrane localization during TGFβ-induced changes in membrane curvature.

### Protein extraction and Western blot analysis

A total of 300,000 cells were seeded per well in 6-well plates and cultured for 24 h before replacing the medium with DMEM containing 5% FBS. Cells were subsequently treated with 12.5 nM doxorubicin for 24 h. Following treatment, cells were washed twice with ice-cold phosphate-buffered saline (PBS) and lysed in RIPA buffer supplemented with protease and phosphatase inhibitor cocktails. Total protein concentration was determined using the Bradford protein assay. Equal amounts of protein (20 μg per sample) were separated by SDS–PAGE and transferred onto PVDF membranes. Membranes were blocked with 5% non-fat dry milk in TBST and incubated overnight at 4 °C with primary antibodies against E-cadherin (kindly provided by Professor Kostantinos Dimas), caveolin-1, LC3, p62 were purchased from Cell Signaling Technology, Danvers, MA, USA) and GAPDH was kindly provided by Professor Ilias Mylonis. After washing, membranes were incubated with the appropriate horseradish peroxidase-conjugated secondary antibodies for 1 h at room temperature. Immunoreactive bands were detected using enhanced chemiluminescence and visualized using a chemiluminescence imaging system. Band intensities were quantified using ImageJ software (subtracting background values) and normalized to GAPDH. The relative protein expression levels were subsequently expressed as fold change relative to the untreated control group.

### Statistical analysis

All experiments were performed independently at least three times (n = 3). Data are presented as the mean ± standard deviation (SD). Statistical analyses were performed using GraphPad Prism. Comparisons between two groups were conducted using an unpaired two-tailed Student’s t-test, whereas comparisons among multiple groups were analyzed using one-way ANOVA followed by Tukey’s multiple-comparison test. Differences were considered statistically significant at P < 0.05.

## Results

### Low-dose doxorubicin induces Caveolin-1 expression in MDA-MB-231 cells

To investigate whether low-dose doxorubicin modulates Caveolin-1 expression in highly invasive breast cancer cells, MDA-MB-231 cells were exposed doxorubicin, and Caveolin-1 protein levels were assessed ((Fig. 1A). Low-dose doxorubicin treatment resulted in a 2.5-fold increase in Caveolin-1 protein expression compared with untreated control cells (P < 0.01; Figure 1B).

**Figure 1.**
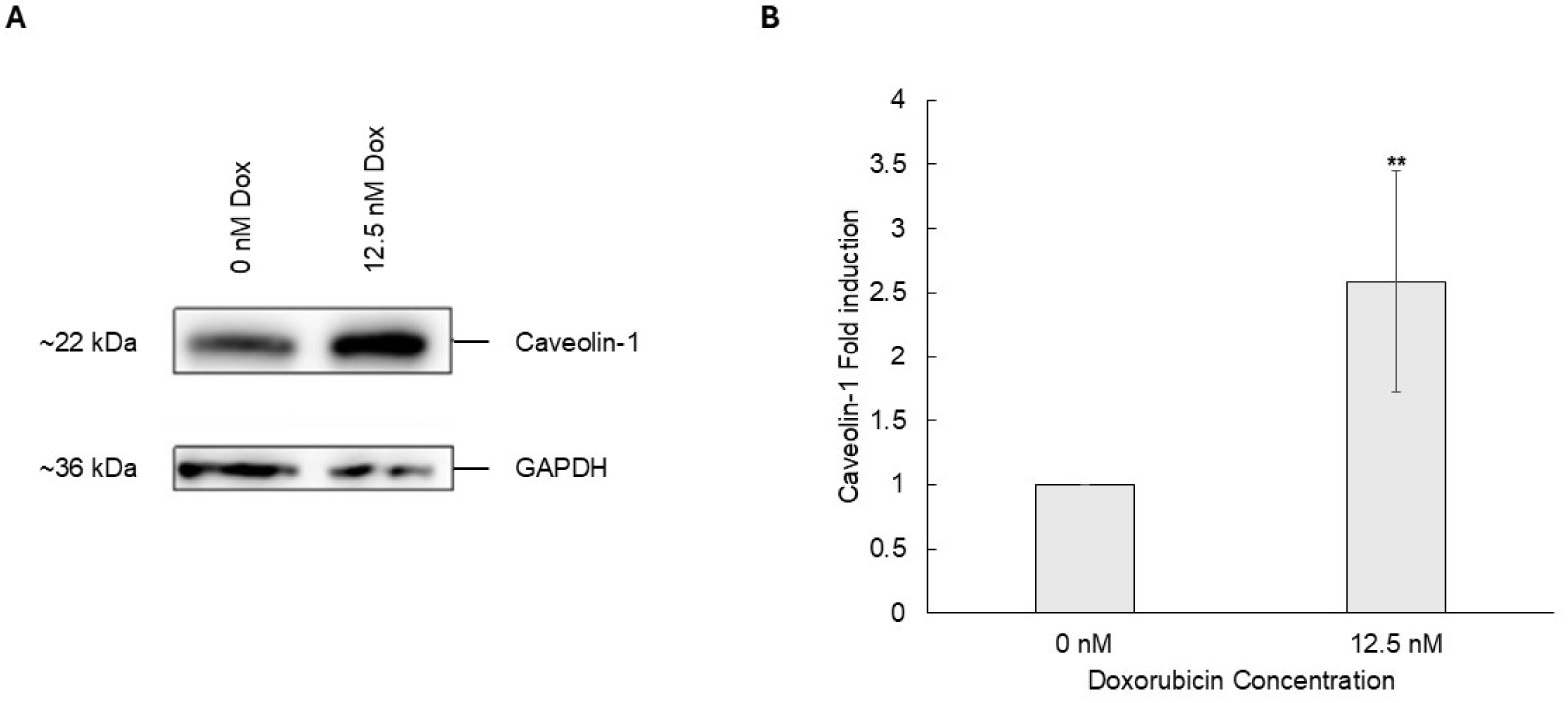
Low-dose doxorubicin increases Caveolin-1 protein expression in MDA-MB-231 cells. **(A)** Representative Western blot of Caveolin-1 expression in untreated (Control) and 12.5 nM doxorubicin-treated MDA-MB-231 cells. GAPDH was used as the loading control. **(B)** Densitometric analysis of Caveolin-1 protein levels normalized to GAPDH and expressed as fold change relative to the untreated control. Bars represent the mean ± SD from five independent experiments (n=5). Statistical significance was determined using an unpaired two-tailed Student’s t-test. **P < 0.01 compared to the control experimental group.

Caveolin-1 is a principal structural component of caveolae and plays a critical role in regulating plasma membrane organization, membrane curvature, mechanotransduction, and compartmentalization of intracellular signaling pathways. Therefore, the observed induction of Caveolin-1 suggests that low-dose doxorubicin initiates an adaptive cellular response involving remodeling of the plasma membrane architecture rather than promoting extensive cytotoxic damage. Increased Caveolin-1 expression may alter the organization of membrane microdomains, thereby influencing the localization and activity of signaling molecules associated with the plasma membrane.

Since Caveolin-1 has been implicated in the regulation of epithelial plasticity, membrane trafficking, and oncogenic signaling, its upregulation provides a mechanistic basis for the subsequent experiments examining epithelial marker expression, Ras membrane localization, and autophagy.

### Low-dose doxorubicin enhances E-cadherin expression and promotes epithelial-like morphological changes in MDA-MB-231 cells

Treatment of MDA-MB-231 cells with doxorubicin resulted in substantial changes in a 1.7-fold increase in E-cadherin protein expression following changes in cellular morphology. (P < 0.05; Fig. 2A, B).

**Figure 2.**
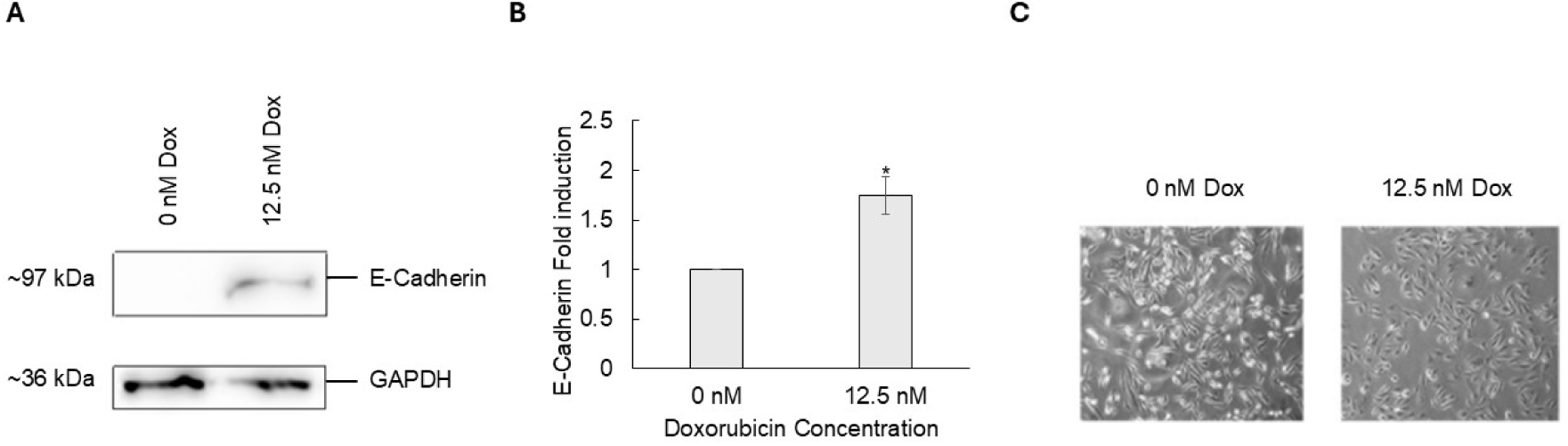
Low-dose doxorubicin enhances E-cadherin expression and promotes epithelial-like morphological changes in MDA-MB-231 cells after 24 h of treatment. **(A)** Representative Western blot of E-cadherin protein expression in untreated control cells and cells treated with 12.5 nM doxorubicin for 24 h. **(B)** Densitometric analysis of E-cadherin protein levels normalized to GAPDH and expressed as fold change relative to the control experimental group. Data are presented as mean ± SD from three independent experiments (n = 3). Statistical significance was determined relative to the control group (*P < 0.05). **(C)** Representative phase-contrast microscopy images of untreated and 12.5 nM doxorubicin-treated MDA-MB-231 cells following 24 h of treatment. Scale bar, 100 μm.

Phase-contrast microscopy was used to examine the morphology of MDA-MB-231 cells after 24 h of treatment. Untreated control cells predominantly exhibited an elongated, spindle-shaped morphology with relatively dispersed growth. In contrast, cells exposed to 12.5 nM doxorubicin displayed a more compact cellular arrangement with increased cell-cell contacts and a cobblestone-like appearance. The overall cell population showed reduced elongation compared with untreated cells while maintaining normal cellular integrity throughout the treatment period. Representative images illustrating these morphological characteristics are shown in Figure 2C.

Therefore, the biochemical and microscopic analyses document increased E-cadherin protein levels together with observable changes in cell morphology following 24 h of exposure to 12.5 nM doxorubicin.

### Doxorubicin induces cytoplasmic redistribution of lipid-anchored tH

To determine whether low-dose doxorubicin alters the subcellular localization of membrane-associated Ras lipid anchors, MDA-MB-231 cells transiently expressing GFP-tagged membrane-targeted tH were treated with doxorubicin for 24 h and analyzed by quantitative fluorescence microscopy. Representative fluorescence images revealed a pronounced difference in tH localization between untreated and doxorubicin-treated cells (Fig. 3A). In control cells, GFP-tH fluorescence was predominantly enriched at the cell periphery, consistent with localization at the plasma membrane. In contrast, cells exposed to 12.5 nM doxorubicin displayed reduced peripheral fluorescence and a more diffuse intracellular distribution, indicating decreased plasma membrane association and increased cytoplasmic localization of tH.

**Figure 3.**
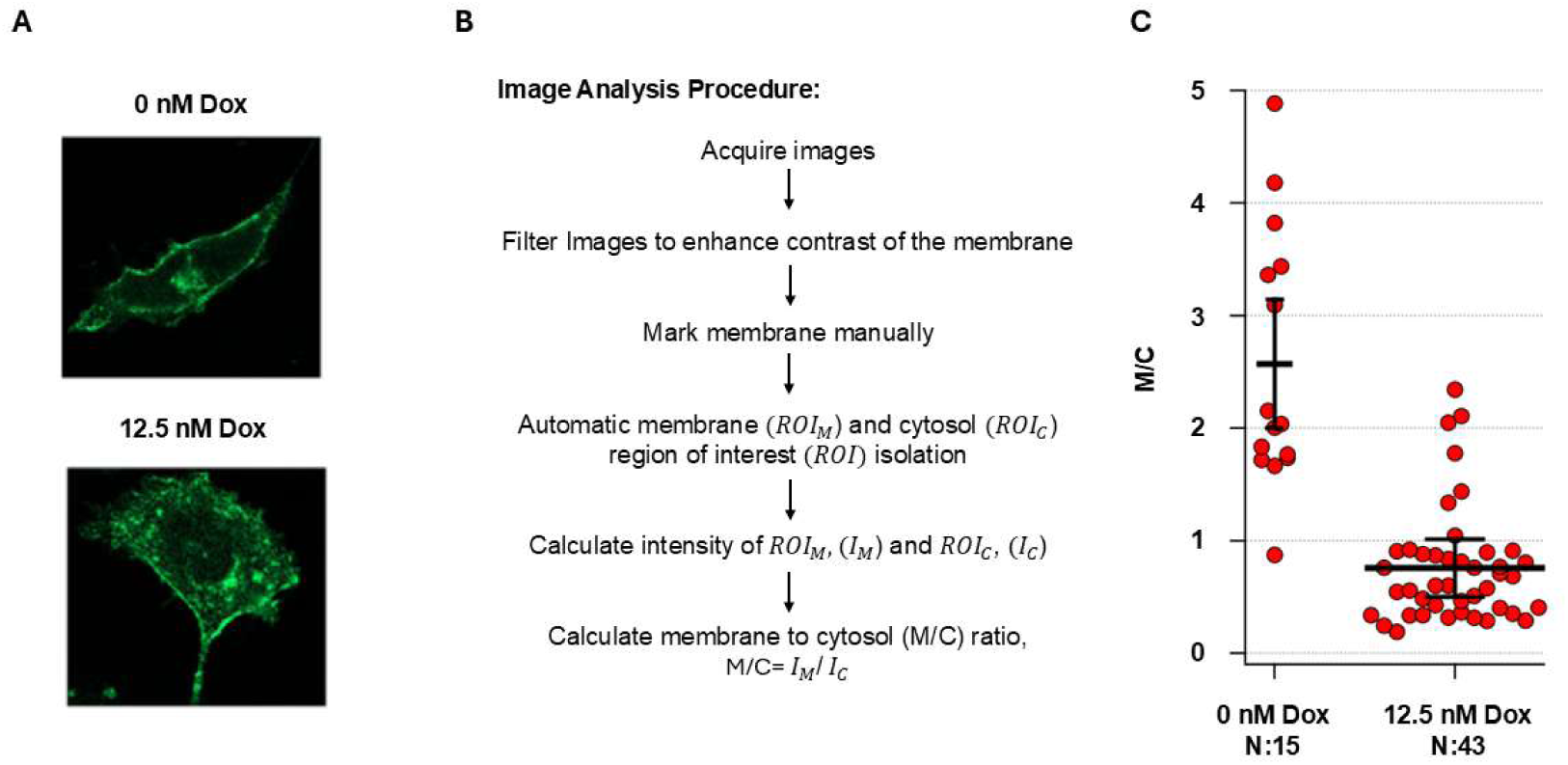
Low-dose doxorubicin induces cytoplasmic redistribution of the H-Ras lipid anchor tH in MDA-MB-231 cells. **(A)** Representative fluorescence images of MDA-MB-231 cells transiently expressing GFP-tH under control conditions or following treatment with 12.5 nM doxorubicin (Dox) for 24 h.. **(B)** Schematic representation of the image-analysis procedure used to quantify GFP-tH localization. Following image acquisition and filtering to enhance membrane contrast, the plasma membrane was manually marked, and membrane (ROI_M) and cytosolic (ROI_C) regions of interest were automatically isolated. Fluorescence intensities within the membrane (I_M) and cytosolic (I_C) regions were measured, and the membrane-to-cytosol ratio was determined following the mathematic formula; M/C = I_M/I_C. **(C)** Assessment of the GFP-tH membrane-to-cytosol (M/C) fluorescence intensity ratio in control (N = 15 cells) and 12.5 nM doxorubicin-treated cells (N = 43 cells). The reduction in the M/C ratio was highly significant (p = 2.04 × 10⁻⁵). Individual points represent individual cells; horizontal and vertical bars indicate the mean and SD, respectively.

To quantitatively assess this redistribution, fluorescence images were analyzed using a membrane-to-cytosol (M/C) intensity-based approach (Figure 3B).

Quantitative analysis demonstrated a marked reduction in the M/C ratio following doxorubicin treatment (N = 43 analyzed cells) compared with untreated control cells (N = 15 analyzed cells) (p = 2.04 × 10⁻⁵; Figure 3C). Control cells exhibited relatively high M/C values, consistent with preferential localization of tH at the plasma membrane, while doxorubicin treated MDA-MB-231 cells showed decreased membrane-associated fluorescence and increased cytoplasmic distribution.

Together, the representative fluorescence images, image-analysis procedure, and quantitative measurements demonstrate that low-dose doxorubicin significantly alters the subcellular distribution of lipid-anchored tH. Specifically, treatment with 12.5 nM doxorubicin for 24 h reduces the relative association of tH with the plasma membrane and promotes its redistribution toward the cytoplasm, resulting in a pronounced decrease in the membrane-to-cytosol fluorescence ratio. These findings establish a measurable effect of low-dose doxorubicin on tH subcellular localization.

### Low-dose doxorubicin promotes accumulation of autophagy-associated proteins

Treatment of MDA-MB-231 breast cancer cells with 12.5 nM doxorubicin for 24 h resulted in marked alterations in the expression of proteins involved in the autophagy pathway (Fig. 4). More specifically, Western blot analysis demonstrated changes in the levels of LC3, ATG5, and p62 compared with untreated control cells, indicating that low-dose doxorubicin substantially affected the autophagic machinery, as it is shown in Figure 4A, while quantitatve densitometric analyses from three independent experiments are shown in Figure 4B–E. The coordinated activities of ATG5, LC3 processing, and p62 turnover determine autophagic flux (Mizushima, 2025). While increases in LC3-II may indicate enhanced autophagosome formation, concomitant accumulation of p62 frequently reflects impaired autophagic degradation rather than efficient completion of the pathway (Runwal et al., 2019).

**Figure 4.**
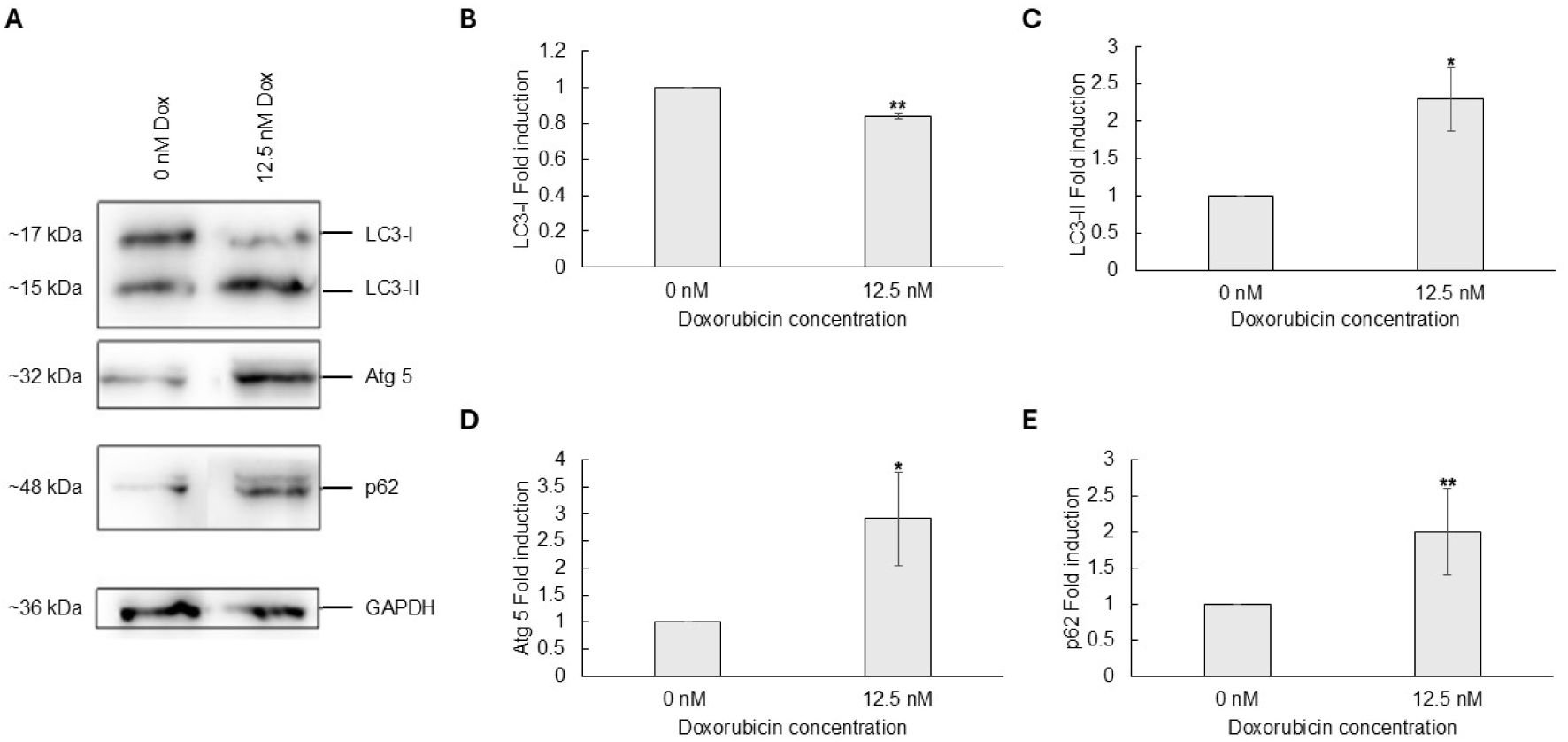
Doxorubicin treatment induces accumulation of autophagy-related proteins in MDA-MB-231 cells. **(A)** Representative Western blots of the autophagy markers LC3, ATG5, p62, and GAPDH in untreated control cells and the respective cells treated with 12.5 nM doxorubicin for 24 h. LC3-I and LC3-II bands are indicated. **(B–E)** Densitometric analysis of (B) LC3-I, (C) LC3-II, (D) ATG5, and **(E)** p62 protein expression normalized to GAPDH and expressed as fold change relative to untreated controls.. Data are presented as mean ± SD from three independent experiments (n = 3). Statistical significance was determined using an unpaired two-tailed Student’s t-test. *P < 0.05, **P < 0.01 versus control.

**Figure 5.**
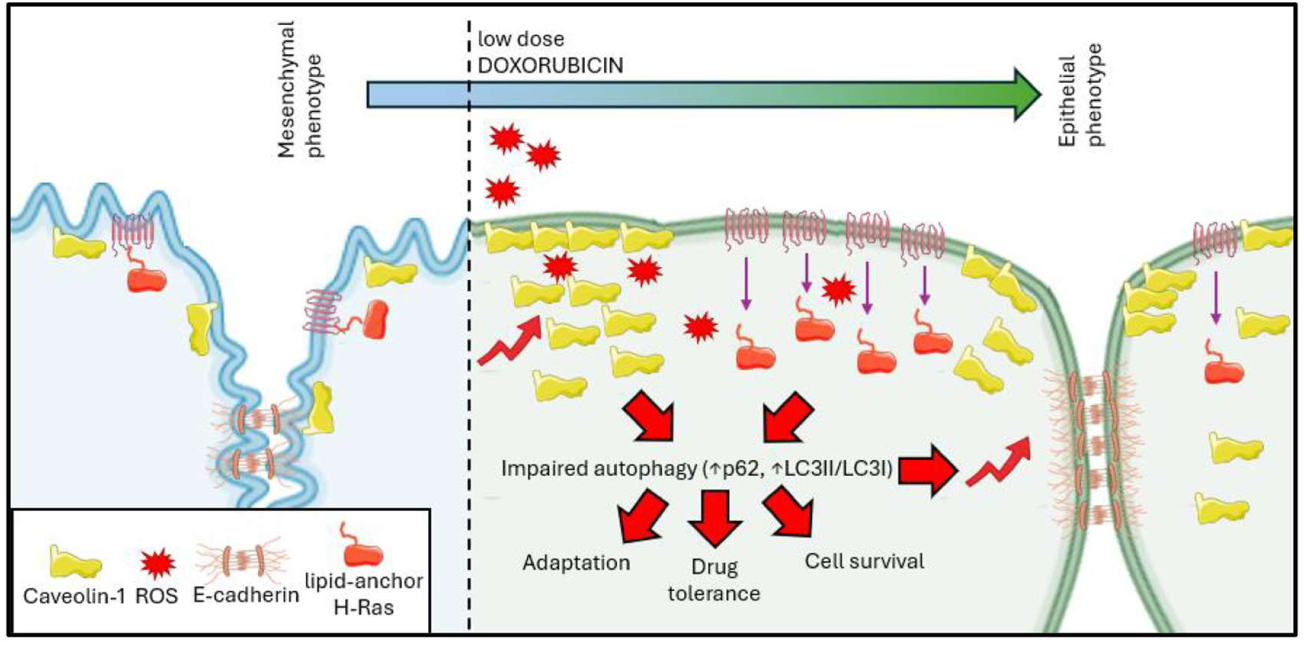
Proposed mechanism underlying low-dose doxorubicin-induced phenotypic remodeling in MDA-MB-231 cells. Schematic representation of the proposed mechanism by which low-dose doxorubicin promotes phenotypic and cellular adaptation in MDA-MB-231 cells. Treatment with low-dose doxorubicin induces reactive oxygen species (ROS), accompanied by increased Caveolin-1 expression. Elevated Caveolin-1 is associated with increased E-cadherin expression and the transition from a mesenchymal toward a more epithelial phenotype. Concomitantly, remodeling of the plasma membrane environment promotes redistribution of the H-Ras lipid anchor (tH) from the plasma membrane toward the cytoplasm. Low-dose doxorubicin is also associated with impaired autophagy, characterized by increased p62 accumulation and an elevated LC3-II/LC3-I ratio. Collectively, these coordinated changes are proposed to represent an adaptive cellular response to low-dose doxorubicin that may contribute to drug tolerance and cell survival.

LC3-I protein levels revealed a statistical significant reduction (**p<0,01) following doxorubicin treatment relative to untreated cells (Figure 4B). In contrast, LC3-II expression was significantly increased after treatment (Figure 4C), demonstrating enhanced accumulation of the lipidated form of LC3. The reciprocal decrease in LC3-I together with the increase in LC3-II (p value) is consistent with increased formation and retention of autophagosomal membranes.

To further assess the observed result, we analysed ATG5 expression. In our experimental setup we have found an increase in ATG5 protein abundance in doxorubicin-treated cells compared to the respective control treated cells (Figure 4D). As ATG5 is an essential component of the autophagosome formation machinery, its elevated expression indicates activation of the molecular components required for autophagosome biogenesis.

Similarly, p62/SQSTM1 protein levels were markedly elevated in response to low-dose doxorubicin (Figure 4E). Because p62 functions as an autophagy adaptor that is normally degraded during efficient autophagic turnover, its accumulation together with increased LC3-II suggests that autophagosomes accumulate without being efficiently degraded. Thus, the simultaneous elevation of LC3-II and p62 provides evidence that doxorubicin treatment is associated with impaired autophagic flux rather than complete autophagic degradation.

Overall, treatment of MDA-MB-231 cells with 12.5 nM doxorubicin for 24 h induced changes in multiple autophagy-related proteins. promoting autophagosome accumulation but impaired autophagy degradation in MDA-MB-231 cells under the experimental conditions used in this study.

## Discussion

The present study provides evidence that low-dose doxorubicin elicits a coordinated adaptive response in MDA-MB-231 breast cancer cells that extends beyond classical cytotoxic activity. By combining biochemical analysis, live-cell imaging, and morphological assessment, we demonstrate that treatment with 12.5 nM doxorubicin increases caveolin-1 expression, enhances E-cadherin levels, redistributes the membrane-targeting H-Ras lipid anchor from the plasma membrane to the cytoplasm, promoting accumulation of LC3-II, ATG5, and p62. Collectively, these findings support a model in which plasma membrane remodeling constitutes a central event linking epithelial plasticity with altered Ras membrane organization and dysregulated autophagy.

One of the major observations is the robust induction of caveolin-1 following exposure to low-dose doxorubicin. This effect is attributed to doxorubicin induced ROS production (Kim et al., 2006) which is associated with caveolin 1 upregulation expression. (Goutas et al., 2020). Caveolin-1 is the principal structural protein of caveolae and is a critical determinant of membrane curvature, cholesterol homeostasis, mechano-transduction, and compartmentalization of signaling molecules. Previous studies have established that caveolin-1 exhibits context-dependent functions during cancer progression (Segal et al., 2025). In some tumors its loss facilitates transformation, whereas in others increased expression promotes cellular adaptation to stress (Simon et al., 2020). Our findings suggest that restoration of caveolin-1 represents an adaptive remodeling response that accompanies epithelial reprogramming rather than generalized toxicity. Increased caveolin-1 may stabilize membrane microdomains, modify membrane mechanics, and reorganize signaling platforms that regulate proliferation and migration.

The increase in E-cadherin together with the transition from elongated spindle-shaped cells toward a compact epithelial-like morphology strongly supports partial mesenchymal-to-epithelial transition. Complete MET was not examined; however, restoration of E-cadherin is widely recognized as an early hallmark of epithelial differentiation. Emerging evidence indicates that chemotherapy can induce phenotypic plasticity depending on dose, duration and cellular context (Iborra et al., 2023). Our observations therefore suggest that sub-cytotoxic doxorubicin concentrations remodel cellular identity while preserving viability, potentially creating an intermediate epithelial state.

A particularly novel aspect of this work is the quantitative demonstration that low-dose doxorubicin alters the intracellular localization of the lipid-anchor H-Ras membrane-targeting sequence. The tH probe faithfully reports plasma membrane association because its localization depends on lipid composition and membrane biophysical properties (Sezgin et al., 2017) (Larsen et al., 2017) (Larsen et al., 2015) (Zhou and Hancock, 2023) (Liang et al., 2019). The marked reduction in membrane-to-cytosol ratio indicates that membrane remodeling decreases the affinity of the lipid anchor for the plasma membrane. This observation is consistent with our previous work proposing that epithelial reprogramming and membrane curvature influence Ras membrane organization (Damalas et al., 2022) (Yang et al., 2024) (Damalas et al., 2026) (Kuburich et al., 2023). Although the present experiments used the isolated lipid anchor rather than full-length H-Ras, the results strongly support the concept that changes in membrane architecture directly influence membrane targeting of Ras proteins.

The autophagy data provide additional mechanistic insight. Increased ATG5 expression suggests activation of the molecular machinery responsible for autophagosome formation, whereas accumulation of LC3-II indicates expansion or persistence of autophagosomal membranes (Ye et al., 2018). Importantly, simultaneous accumulation of p62 argues against efficient autophagic degradation and instead supports impaired autophagic flux (Quan et al., 2025). This distinction is biologically important because increased LC3-II alone cannot discriminate between enhanced autophagy initiation and defective lysosomal clearance. The combined elevation of LC3-II and p62 therefore indicates that autophagosomes accumulate without efficient degradation.

Integration of all experimental observations allows the proposal of a unified mechanistic model. Low-dose doxorubicin likely induces oxidative and membrane stress, resulting in remodeling of plasma membrane organization and increased caveolin-1 expression. Changes in membrane curvature reduce plasma membrane association of the H-Ras lipid anchor, while simultaneous induction of E-cadherin promotes epithelial reorganization. In parallel, the concurrent increase in caveolin-1, LC3-II and p62 is indicating an impaired autophagic clearance, as caveolin-1 has been reported to negatively regulate autophagy by limiting autophagosome–lysosome fusion in breast cancer cells (Shi et al., 2015). Importantly, previous studies from our lab demonstrated that siRNA mediated depletion of caveolin 1 enhances autophagy (Goutas et al., 2021). Rather than independent phenomena, these events appear to represent interconnected components of a coordinated adaptive response.

The findings are particularly relevant because membrane organization has emerged as a major regulator of oncogenic signaling. Ras proteins require precise nanoscale membrane organization for efficient signal transduction, and alterations in lipid composition, cholesterol content or membrane curvature profoundly influence Ras nanoclustering and downstream signaling (Damalas et al., 2022) (Damalas et al., 2026) (Liang et al., 2019) (Zhou and Hancock, 2023) (Larsen et al.). Consequently, therapeutic strategies capable of modifying membrane architecture may indirectly regulate oncogenic pathways independently of direct kinase inhibition. Our results provide experimental support for this concept by demonstrating that pharmacological treatment alters the localization of a Ras membrane-targeting probe.

To our knowledge, very limited studies have simultaneously examined caveolin-1 induction, redistribution of a lipid-anchor H-Ras biosensor and defective autophagic flux within the same experimental framework. This integrated approach provides a broader understanding of how membrane biophysics contributes to cellular adaptation during chemotherapy.

In conclusion, our data support a model in which low-dose doxorubicin promotes membrane remodeling characterized by caveolin-1 induction, increased epithelial differentiation, altered H-Ras membrane association and impaired autophagic flux. These coordinated responses identify plasma membrane architecture as an active regulator of tumor cell plasticity rather than a passive structural element. Understanding how membrane organization integrates Ras signaling, epithelial reprogramming and autophagy may facilitate development of therapeutic strategies that target adaptive responses responsible for tumor persistence and treatment resistance.

## Author Contributions

A.D. conceived and designed the research, conducted the experiments, and wrote the original draft of the manuscript. M.T. and E.V. performed the data analysis and data curation. I.D.K. reviewed, edited the manuscript and involved in the preparation of the illustration. P.M.D. performed the statistical analyses and involved in the preparation of the figures. S.K, Z.O, A.C, and P.L. reviewed and edited the manuscript. B.T. supervised the overall study and contributed to the writing and editing of the manuscript. All authors contributed to the article and approved the submitted version.

## Acknowledgments

We would like to thank Professor Daniel Abankwa for providing the GFP-tH plasmid, generously. We gratefully acknowledge Cristina Rosi Vasiliou for her valuable contribution to this work. The research is conducted in the operating framework of the University of Thessaly Innovation, Technology Transfer Unit and Entrepreneurship Center “One Planet Thessaly”, under the “University of Thessaly Grants for Scientific Publication Support” action and is funded by the Special Account of Research Grants of the University of Thessaly. This research was also funded by the Research Council of Lithuania, grant number S-MIP-24-58 (to M.T.).

## References

Abankwa, D. & Gorfe, A. A. 2020. Mechanisms of Ras Membrane Organization and Signaling: Ras Rocks Again. Biomolecules, 10.

Allgayer, H., Mahapatra, S., Mishra, B., Swain, B., Saha, S., Khanra, S., Kumari, K., Panda, V. K., Malhotra, D., Patil, N. S., Leupold, J. H. & Kundu, G. C. 2025. Epithelial-to-mesenchymal transition (EMT) and cancer metastasis: the status quo of methods and experimental models 2025. Mol Cancer, 24, 167.

Ariotti, N., Fernandez-Rojo, M. A., Zhou, Y., Hill, M. M., Rodkey, T. L., Inder, K., Tanner, L. B., Wenk, M. R., Hancock, J. F. & Parton, R. G. 2014. Caveolae regulate the nanoscale organization of the plasma membrane to remotely control Ras signaling. Journal of Cell Biology, 204, 777.

Ariotti, N., Rae, J., Leneva, N., Ferguson, C., Loo, D., Okano, S., Hill, M. M., Walser, P., Collins, B. M. & Parton, R. G. 2015. Molecular Characterization of Caveolin-induced Membrane Curvature. J Biol Chem, 290, 24875–90.

Bruner, H. C. & Derksen, P. W. B. 2018. Loss of E-Cadherin-Dependent Cell-Cell Adhesion and the Development and Progression of Cancer. Cold Spring Harb Perspect Biol, 10.

Chan, T. S., Hsu, C. C., Pai, V. C., Liao, W. Y., Huang, S. S., Tan, K. T., Yen, C. J., Hsu, S. C., Chen, W. Y., Shan, Y. S., Li, C. R., Lee, M. T., Jiang, K. Y., Chu, J. M., Lien, G. S., Weaver, V. M. & Tsai, K. K. 2016. Metronomic chemotherapy prevents therapy-induced stromal activation and induction of tumor-initiating cells. J Exp Med, 213, 2967–2988.

Coopman, P. & Djiane, A. 2016. Adherens Junction and E-Cadherin complex regulation by epithelial polarity. Cell Mol Life Sci, 73, 3535–53.

Damalas, A., Kyriazis, I. D., Tutkus, M., Angelidis, C. & Trachana, V. 2026. Membrane Curvature and Cancer: Mechanisms, Implications, and Therapeutic Perspectives. Cancers (Basel), 18.

Damalas, A., Vonkova, I., Tutkus, M. & Stamou, D. 2022. TGFbeta-induced changes in membrane curvature influence Ras oncoprotein membrane localization. Sci Rep, 12, 13486.

Datta, A., Deng, S., Gopal, V., Yap, K. C., Halim, C. E., Lye, M. L., Ong, M. S., Tan, T. Z., Sethi, G., Hooi, S. C., Kumar, A. P. & Yap, C. T. 2021. Cytoskeletal Dynamics in Epithelial-Mesenchymal Transition: Insights into Therapeutic Targets for Cancer Metastasis. Cancers (Basel), 13.

Dhiman, V. K., Kumari, M. & Singh, D. 2026. Chemoresistance: The hidden barrier in cancer treatment. Cancer Pathog Ther, 4, 98–109.

Filippini, A. & D’alessio, A. 2020. Caveolae and Lipid Rafts in Endothelium: Valuable Organelles for Multiple Functions. Biomolecules, 10.

Goutas, A., Outskouni, Z., Papathanasiou, I., Satra, M., Koliakos, G. & Trachana, V. 2021. Dysregulation of Caveolin-1 Phosphorylation and Nuclear Translocation Is Associated with Senescence Onset. Cells, 10.

Goutas, A., Papathanasiou, I., Mourmoura, E., Tsesmelis, K., Tsezou, A. & Trachana, V. 2020. Oxidative Stress Response Is Mediated by Overexpression and Spatiotemporal Regulation of Caveolin-1. Antioxidants (Basel), 9.

Iborra, F. J., Marti, C., Calabuig-Navarro, V., Papadopoulos, P., Meseguer, S., Iborra, P. M., Garcia, F., Martinez-Lorente, A., Almazan, F. & Calabuig, J. 2023. Chemotherapy induces cell plasticity; controlling plasticity increases therapeutic response. Signal Transduct Target Ther, 8, 256.

Johnson-Arbor, K., Douedi, S. & Patel, P. 2026. Doxorubicin. StatPearls. Treasure Island (Fl) ineligible companies. Disclosure: Steven Douedi declares no relevant financial relationships with ineligible companies. Disclosure: Preeti Patel declares no relevant financial relationships with ineligible companies.

Katsuno-Kambe, H., Parton, R. G., Yap, A. S. & Teo, J. L. 2021. Caveolin-1 influences epithelial collective cell migration via FMNL2 formin. Biol Cell, 113, 107–117.

Kciuk, M., Gielecinska, A., Mujwar, S., Kolat, D., Kaluzinska-Kolat, Z., Celik, I. & Kontek, R. 2023. Doxorubicin-An Agent with Multiple Mechanisms of Anticancer Activity. Cells, 12.

Kim, S. Y., Kim, S. J., Kim, B. J., Rah, S. Y., Chung, S. M., Im, M. J. & Kim, U. H. 2006. Doxorubicin-induced reactive oxygen species generation and intracellular Ca2+ increase are reciprocally modulated in rat cardiomyocytes. Exp Mol Med, 38, 535–45.

Kuburich, N. A., Sabapathy, T., Demestichas, B. R., Maddela, J. J., Den Hollander, P. & Mani, S. A. 2023. Proactive and reactive roles of Tgf-beta in cancer. Semin Cancer Biol, 95, 120–139.

Lamouille, S., Xu, J. & Derynck, R. 2014. Molecular mechanisms of epithelial-mesenchymal transition. Nat Rev Mol Cell Biol, 15, 178–96.

Larsen, J. B., Jensen, M. B., Bhatia, V. K., Pedersen, S. L., Bjornholm, T., Iversen, L., Uline, M. J., Szleifer, I., Jensen, K. J., Hatzakis, N. S. & Stamou, D. 2015. Membrane curvature enables N-Ras lipid anchor sorting to liquid-ordered membrane phases. Nature Chemical Biology, 11, 192–U176.

Larsen, J. B., Kennard, C., Pedersen, S. L., Jensen, K. J., Uline, M. J., Hatzakis, N. S. & Stamou, D. 2017. Membrane Curvature and Lipid Composition Synergize To Regulate N-Ras Anchor Recruitment. Biophys J, 113, 1269–1279.

Lemanowicz, J., Kloska, S. M., Siwik-Ziomek, A., Kolaczyk, P., Lipinska, U. W. & Kloska, A. 2026. Biochemical Mechanisms of Cellular Stress Adaptation in the Pathogenesis of Chronic Diseases. Molecules, 31.

Liang, H., Mu, H., Jean-Francois, F., Lakshman, B., Sarkar-Banerjee, S., Zhuang, Y., Zeng, Y., Gao, W., Zaske, A. M., Nissley, D. V., Gorfe, A. A., Zhao, W. & Zhou, Y. 2019. Membrane curvature sensing of the lipid-anchored K-Ras small GTPase. Life Sci Alliance, 2.

Liu, S., Dong, Y., Wang, Y., Hu, P., Wang, J. & Wang, R. Y. 2021. Pristimerin exerts antitumor activity against Mda-Mb-231 triple-negative breast cancer cells by reversing of epithelial-mesenchymal transition via downregulation of integrin beta3. Biomed J, 44, S84–S92.

Lukasiewicz, S., Czeczelewski, M., Forma, A., Baj, J., Sitarz, R. & Stanislawek, A. 2021. Breast Cancer-Epidemiology, Risk Factors, Classification, Prognostic Markers, and Current Treatment Strategies-An Updated Review. Cancers (Basel), 13.

Luo, M., Brooks, M. & Wicha, M. S. 2015. Epithelial-mesenchymal plasticity of breast cancer stem cells: implications for metastasis and therapeutic resistance. Curr Pharm Des, 21, 1301–10.

Mizushima, N. 2025. Autophagic flux measurement: Cargo degradation versus generation of degradation products. Curr Opin Cell Biol, 93, 102463.

Newton, E. E., Mueller, L. E., Treadwell, S. M., Morris, C. A. & Machado, H. L. 2022. Molecular Targets of Triple-Negative Breast Cancer: Where Do We Stand? Cancers (Basel), 14.

Nicolini, A., Ferrari, P., Silvestri, R. & Bini, D. A. 2025. Targeted Therapies in the Most Common Advanced Solid Tumors, Drug Resistance, and Counteracting Progressive Micrometastatic Disease: The Next Frontier of Research. MedComm (2020), 6, e70373.

Parton, R. G. & Simons, K. 2007. The multiple faces of caveolae. Nat Rev Mol Cell Biol, 8, 185–94.

Pilco-Ferreto, N. & Calaf, G. M. 2016. Influence of doxorubicin on apoptosis and oxidative stress in breast cancer cell lines. Int J Oncol, 49, 753–62.

Quan, X., Yang, Y., Liu, X., Kaltwasser, B., Pillath-Eilers, M., Walkenfort, B., Voortmann, S., Mohamud Yusuf, A., Hagemann, N., Wang, C., Hasenberg, M., Hermann, D. M. & Brockmeier, U. 2025. Autophagy hub-protein p62 orchestrates oxidative, endoplasmic reticulum stress, and inflammatory responses post-ischemia, exacerbating stroke outcome. Redox Biol, 84, 103700.

Quest, A. F., Gutierrez-Pajares, J. L. & Torres, V. A. 2008. Caveolin-1: an ambiguous partner in cell signalling and cancer. J Cell Mol Med, 12, 1130–50.

Runwal, G., Stamatakou, E., Siddiqi, F. H., Puri, C., Zhu, Y. & Rubinsztein, D. C. 2019. LC3-positive structures are prominent in autophagy-deficient cells. Sci Rep, 9, 10147.

Segal, D., Wang, X., Mazloom-Farisbaf, H., Rajendran, D., Butler, E., Chen, B., Chang, B. J., Ahuja, K., Perny, A., Bhatt, K., Reed, D. K., Castrillon, D. H., Lee, J., Jeffery, E., Wang, L., Nguyen, K., Williams, N. S., Skapek, S. X., Rajaram, S., Fiolka, R., Jaqaman, K., Hon, G., Amatruda, J. F. & Danuser, G. 2025. Caveolin-1 regulates context-dependent signaling and survival in Ewing sarcoma. bioRxiv.

Sezgin, E., Levental, I., Mayor, S. & Eggeling, C. 2017. The mystery of membrane organization: composition, regulation and roles of lipid rafts. Nat Rev Mol Cell Biol, 18, 361–374.

Shi, Y., Tan, S. H., Ng, S., Zhou, J., Yang, N. D., Koo, G. B., Mcmahon, K. A., Parton, R. G., Hill, M. M., Del Pozo, M. A., Kim, Y. S. & Shen, H. M. 2015. Critical role of CAV1/caveolin-1 in cell stress responses in human breast cancer cells via modulation of lysosomal function and autophagy. Autophagy, 11, 769–84.

Simon, L., Campos, A., Leyton, L. & Quest, A. F. G. 2020. Caveolin-1 function at the plasma membrane and in intracellular compartments in cancer. Cancer Metastasis Rev, 39, 435–453.

Yang, Y., Ma, Q., Yang, M., Wei, R., Wang, Z., Jiang, C., Liu, H. & Han, M. 2025. The crosstalk of caveolin-1 and autophagy in different diseases. Front Immunol, 16, 1648757.

Yang, Y., Valencia, L. A., Lu, C. H., Nakamoto, M. L., Tsai, C. T., Liu, C., Yang, H., Zhang, W., Jahed, Z., Lee, W. R., Santoro, F., Liou, J., Wu, J. C. & Cui, B. 2024. Plasma membrane curvature regulates the formation of contacts with the endoplasmic reticulum. Nat Cell Biol, 26, 1878–1891.

Ye, X., Zhou, X. J. & Zhang, H. 2018. Exploring the Role of Autophagy-Related Gene 5 (ATG5) Yields Important Insights Into Autophagy in Autoimmune/Autoinflammatory Diseases. Front Immunol, 9, 2334.

Zhou, Y. & Hancock, J. F. 2018. Deciphering lipid codes: K-Ras as a paradigm. Traffic, 19, 157–165.

Zhou, Y. & Hancock, J. F. 2023. Ras nanoclusters are cell surface transducers that convert extracellular stimuli to intracellular signalling. Febs Lett, 597, 892–908.

Zhou, Y., Prakash, P., Liang, H., Cho, K. J., Gorfe, A. A. & Hancock, J. F. 2017. Lipid-Sorting Specificity Encoded in K-Ras Membrane Anchor Regulates Signal Output. Cell, 168, 239–251 e16.

